# Mechano-bactericidal nanopillars require external forces to effectively kill bacteria

**DOI:** 10.1101/2020.03.27.012153

**Authors:** Amin Valiei, Nicholas Lin, Jean-Francois Bryche, Geoffrey McKay, Michael Canva, Paul G. Charette, Dao Nguyen, Christopher Moraes, Nathalie Tufenkji

## Abstract

Nanopillars are known to mechanically damage bacteria, suggesting a promising strategy for highly-effective anti-bacterial surfaces. However, the mechanisms underlying this phenomena remain unclear, which ultimately limits translational potential towards real-world applications. Using real-time and end-point analysis techniques, we demonstrate that in contrast to expectations, bacteria on multiple “mechano-bactericidal” surfaces remain viable, unless exposed to a moving air-liquid interface which caused considerable cell death. Reasoning that normal forces arising from surface tension may underlie mechano-bactericidal activity, we developed computational and experimental models to estimate, manipulate, and recreate the impact of these forces. Our experiments together demonstrate that nanopillar surfaces alone do not cause cell death, but require a critical level of external force to deform and rupture bacteria. These studies hence provide fundamental physical insight into the mechanisms by which nanopillar surfaces can serve as effective antibacterial strategies, and describe the use-conditions under which such nanotechnological approaches may provide practical value.

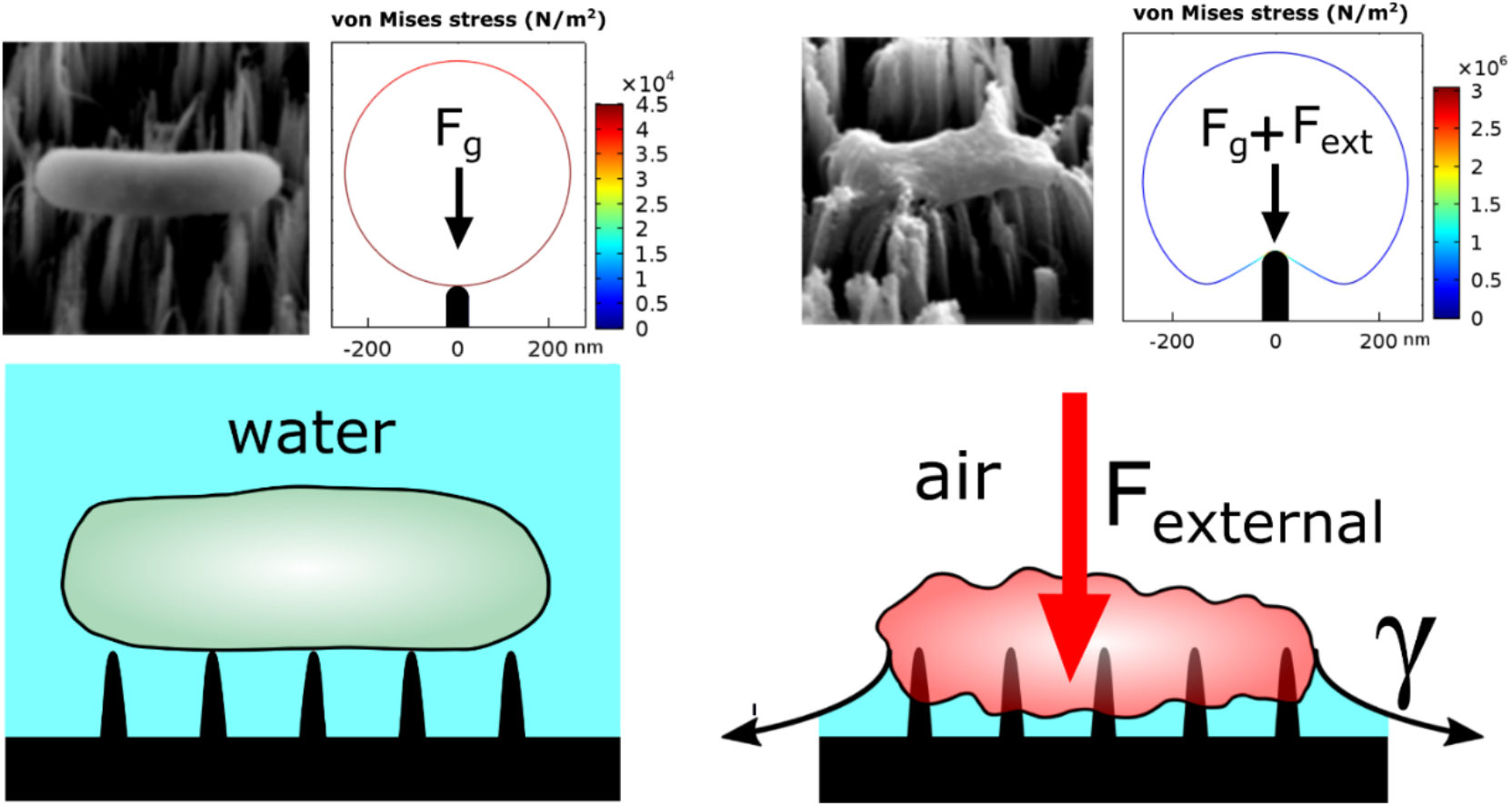
Graphical Abstract.

---

Innovative antimicrobial technologies can play an important role in future designs of surfaces used in hospitals, public transportation, and food production equipment^1–3^. Disinfecting these surfaces through routine application of biocides risks the emergence of antimicrobial-resistant strains^4–5^. Surface immobilization of antibacterial compounds that kill on contact improves cost-effectiveness, but has a limited lifetime as the compounds can degrade; and reduced antibacterial efficacy over time can lead to resistant bacterial strains^6^. To overcome these limitations, antibacterial strategies that rely on physical factors such as the surface’s topography have recently been proposed as a strategy to maintain cost-effective bactericidal efficiency, thereby circumventing evolutionary mechanisms that give rise to antimicrobial resistance.

In 2012, it was reported that the nanopillars on the wings of the *Psaltodo claripennis* cicada insect are capable of structurally deforming bacteria and are therefore considered antibacterial^7^. Changing the surface chemistry of the cicada wing did not alter antibacterial effectiveness, implying that bactericidal activity of natural nanopillar arrays originates from physical interactions that can deform and rupture the bacterial cell wall^8^. Subsequent studies showed similar nanopillar-induced antibacterial behaviour on the wings of other insects, such as damselflies and dragonflies, which have sharper nanopillars said to deliver even higher antibacterial efficacy^9–10^. These initial discoveries prompted the development of engineered nanopillars capable of mimicking this physical antibacterial mechanism such as black silicon nanopillars that mimic nanopillars of dragonfly wings and have remarkable bactericidal activity^11^. Nanopillars constructed from titanium^12^, chitosan^13^, stainless steel^14^, zinc oxide^15^, zinc phthalocyanine^16^, poly(methyl methacrylate)^17^, carbon nanotubes^18^ and others have also shown antibacterial capabilities^19–22^. The diversity of these materials reinforces the notion that the dominant antibacterial mechanism originates from physical structure rather than chemical interactions.

While past works confirm that mechanical forces are responsible for damaging bacteria on nanopillars^20–21^, there is no consensus on the mechanism(s) of action of the killing forces involved. Mechanical damage caused by nanopillars has been attributed by some researchers to adhesion forces which deform cells upon contact with the nanopillars^8, 19^. Alternatively, bacteria may be killed by shear forces generated when attached cells attempt to move laterally on nanopillars^23–24^, or when bending energy stored in high aspect ratio nanopillars is released^25^. Thus, additional clarity is needed regarding the underlying mechanisms of mechano-bactericidal activity, which would ultimately lead to the development of precise theoretical models and practical applications of this technology.

To address this challenge, we studied nanopillar-mediated bacterial killing efficiency of multiple nanopillar surfaces fabricated from nanoparticle-mediated etching of silicon (NanoSi), self-assembly of zinc oxide (NanoZnO), nanolithography of silicon, and natural cicada wings. NanoSi and NanoZnO surfaces were predominantly used as the test surfaces for this study, as both can be controllably fabricated at sufficient throughput to support multiple experiments and conditions. Furthermore, NanoSi is relatively well-studied in the context of mechano-bactericidal surfaces and shows the highest antibacterial performance reported^11, 20^ whereas the hydrothermal synthesis of NanoZnO is well-established in the literature. Morphology of the nanopillars was characterized by scanning electron microscopy (Figure 1). Nanopillars of NanoSi are generally tapered and sharper and narrower than nanopillars of NanoZnO, which are blunt-ended and wider. The dimensions of nanopillars were synthesized to be within the range of other candidate bactericidal surfaces (Table S1), with the pillar diameters spanning the range of previously reported mechano-bactericidal structures (Table 4 in Lin *et al.*^22^ and Table 2 in Tripathy *et al.*^20^). Details of fabrication for NanoSi and NanoZnO as well as fabrication of alternative ordered nanopillar surfaces via nanolithography, and characterization of natural cicada wings is described in the Supporting Information.

**Figure 1.**
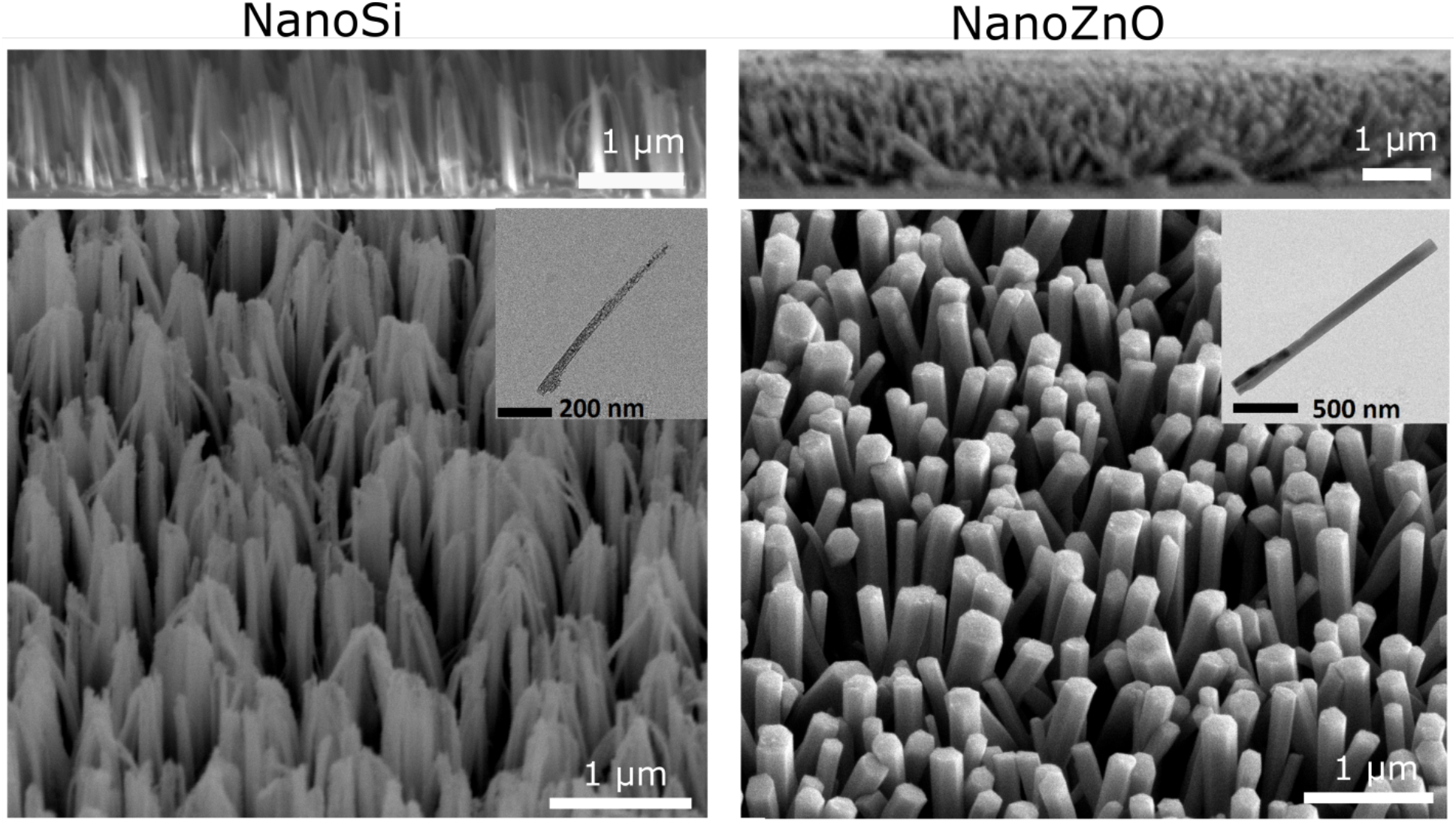
SEM images of NanoSi and NanoZnO surfaces; side view (top), three-dimensional view (bottom). Inset shows a TEM image of one nanopillar. NanoSi consists of sharp vertical high-aspect ratio (diameter = 50 ± 10 nm, height = 0.9 μm) nanopillars and NanoZnO consists of multidirectional nanopillars with blunt-ends (diameter = 230 ± 65 nm and height = 1.2 μm). Nanopillars are randomly distributed on both NanoSi (spacing = 447 ± 395 nm) and NanoZnO (spacing = 408 ± 392 nm).

The model bacterium chosen for this study was *Pseudomonas aeruginosa,* a Gram-negative bacterium known to be a key causative agent of opportunistic human infections^26^. Viability of *P. aeruginosa* was initially evaluated on both NanoSi and NanoZnO nanopillar surfaces while fully immersed in a bacterial suspension labeled with a mixture of live (SYTO 9; green) and dead (propidium iodide; red) fluorescent dyes. Contrary to reports in the literature^11, 15, 27–28^, we detected no antibacterial activity on either nanopillar surface when immersed: fluorescently-labelled bacterial cells adhered to both surfaces immediately but the vast majority of cells (>98%) remained viable after 1 h (Figure 2b and Figure S1).

**Figure 2.**
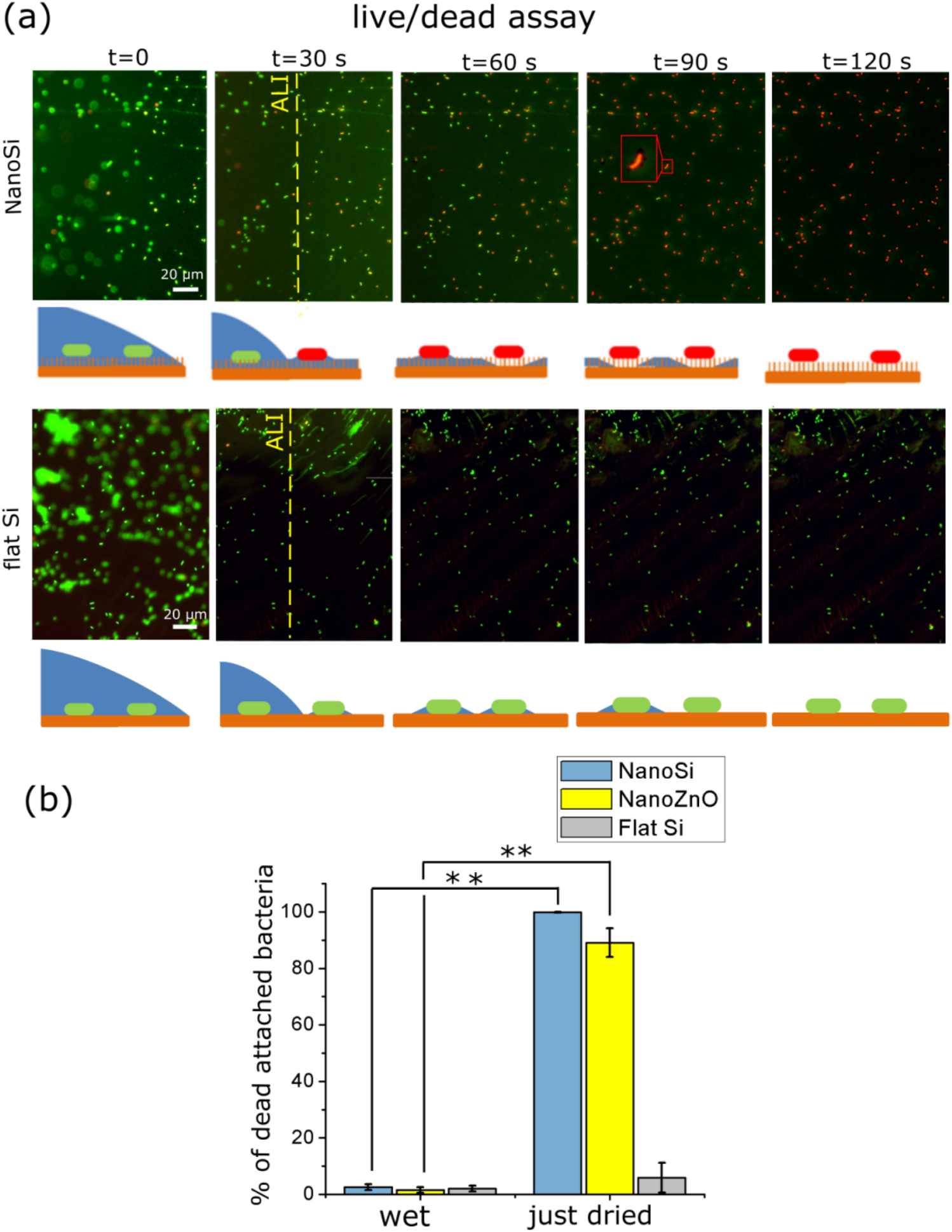
*P. aeruginosa* viability on nanopillars and control (flat) surfaces upon evaporation. (a) viability of bacteria as a function of time on NanoSi and flat Si that is subject to evaporation (live cells are green and dead cells are red). A 3 μL droplet is placed on the surface and the air-liquid interface (ALI) was monitored during the last two minutes of evaporation. At t = 0, the surface is wet. At t = 30 s, the air-liquid interface is passing through the field of view (yellow line shows the approximate position of the ALI). Images at t = 60 s and 90 s show the events that happen just after evaporation (inset at t = 90 s for NanoSi shows the magnified image of a bacterium after the film of water evaporates). At t = 120 s, the surface is completely dry. Sketches below each image graphically depict the possible side view of bacteria deduced from that image (refer to Figure S2 for viability of bacteria on NanoZnO upon evaporation) (b) Percentage of dead cells in the fully immersed wet condition, and immediately after evaporation of water on NanoSi, NanoZnO, and flat Si. Error bars denote the sample standard deviation and the experiments were repeated three times. *Statistically significant difference (*p* < 0.05, n = 3, Student’s t-test).

Although the bacterial suspension was observed to wet the surfaces, similar to previous reports in small-scale experiments^29–30^, we observed randomly entrapped microscopic air bubbles that could be quickly eliminated by gentle agitation. If not eliminated, these air bubbles may spontaneously expand during microscopy, and we observed substantial bacterial killing within the expansion zone (Figure S2), suggesting that the air-liquid interface could play an important role in this process. To better understand how a moving air-liquid interface on a nanopillar surface might affect bacterial viability, a 3 μL droplet of bacterial suspension was placed on the nanopillar surfaces and monitored under fluorescent microscopy. Initially, all attached and planktonic cells observed under the microscope were viable (Figure 2a, t = 0). The droplet volume decreased rapidly due to evaporation which caused the air-liquid interface to recede from the edge towards the droplet center. Monitoring the location where the contact line was passing (represented by yellow line in Figure 2a, t = 30 s), we observed that (i) bacteria were not swept away by the droplet interface but were left behind on the nanopillar surfaces; and (ii) they began to uptake cell membrane-impermeable propidium iodide immediately after air-liquid interface passage (Fig. 2a; t = 60, 90 s), indicating cell death. Over 99.9% of the bacteria on NanoSi (Figure 2a; t=120 s, Figure 2b) and over 90% on NanoZnO (Figure S3, 2b) stained positive for propidium iodide. In contrast, over 95% of bacteria on flat surfaces remained viable even after all liquid evaporated, confirming that this phenomenon is specific to the nanopillar topographies (Fig. 2a,b) and not a result of bacterial dessication or increase in salinity during evaporation. We further confirmed that the results were not an artefact related to the dyes by studying the GFP signal loss upon air-liquid interface displacement with *P. aeruginosa* tagged with green fluorescent protein (GFP) (Figure S4).

The live/dead assay provides information on the integrity of the cell membrane but does not capture bacterial morphologies on the nanopillar surfaces. We therefore used SEM to investigate bacterial structure associated with the live/dead state. As before, a droplet of bacterial suspension was partially dried on each nanopillar surface, and to protect the sample integrity in the just-dried state, the samples were immediately chemically-fixed by glutaraldehyde and prepared for SEM by critical point drying (CPD). CPD retains bacterial morphology during drying by gradually replacing the liquid medium with liquid CO_2_, which is later discharged at supercritical conditions^31^. Carefully following these sample preparation steps allowed us to confirm that bacteria only showed significant cell deformation and damage in the just-dried state on nanopillar surfaces; while bacterial integrity remained intact on control (flat) surfaces or when maintained in fully immersed conditions throughout sample handling (Figure 3, S5). This is consistent with our live/dead assays previously described (Figure 2b, S1), strongly suggesting that that the passage of an air-liquid interface is required to generate appreciable antibacterial activity on the nanopillar surfaces.

**Figure 3.**
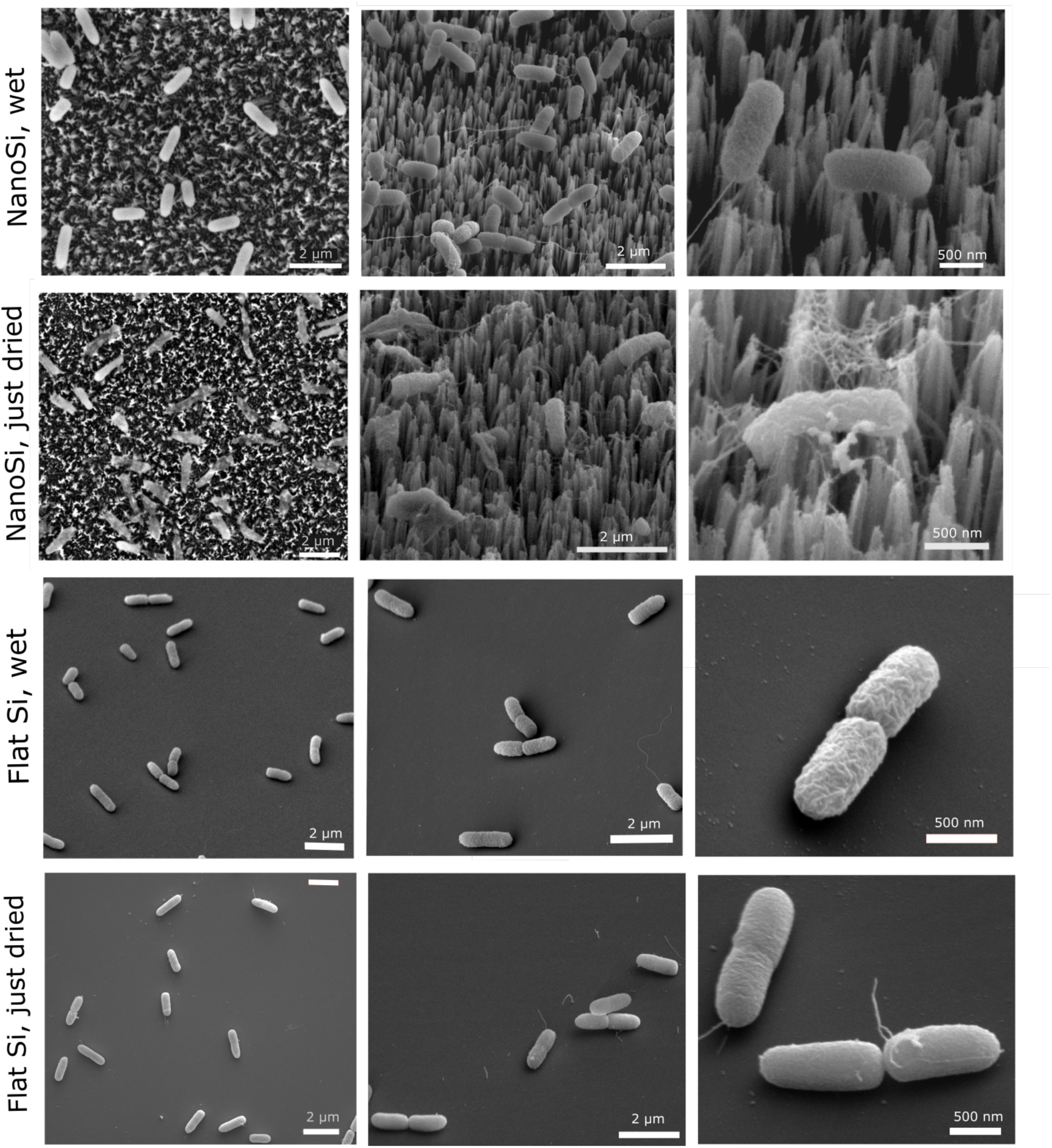
SEM images of *P. aeruginosa* morphology on NanoSi and flat Si at the wet state (samples were not exposed to air, only chemically fixed and critical point-dried) and on NanoSi and flat Si just after water evaporation (samples were air-dried first and after complete water evaporation, chemically fixed and critical point-dried). For SEM images of NanoZnO refer to Figure S3.

To confirm that these surprising findings are not due to the specific NanoSi and NanoZnO platforms used here, we conducted further proof-of-concept studies on other surfaces previously reported to have mechano-bactericidal properties. We studied bacterial morphology on other nanopillar surfaces, including cicada wings of the species *Salvazana mirabilis* and highly ordered silicon nanopillars. Again, SEM observations show cells were damaged on these nanopillar surfaces upon water evaporation, but not in the wet condition (Figure S6), confirming that these results are generalizable across nanopillar surfaces. These results confirm that in contrast with previous studies^11, 15, 27–28^, nanopillar surfaces alone do not exhibit antibacterial effects, unless accompanied by the passage of an air-liquid interface.

Given that the movement of an air-liquid interface is a requirement for the observed bactericidal activity, we reasoned that the substantial normal forces associated with an air-liquid interface^32^ exerted by the capillary action of fluids might be responsible for rupturing bacteria on the nanopillars. To test this hypothesis, we first estimated the magnitude of normal capillary forces during water evaporation using computational approaches, and then experimentally validated this approach by manipulating physical parameters associated with normal forces.

Capillary forces arise when the evaporating liquid level falls below the bacterial cell height (Figure 4a). The first component of the capillary forces is the surface tension force (**F_s_**) which acts around the periphery of the meniscus along the liquid/vapor interface^33–34^. **F_s_** is calculated using:

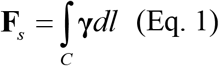

**Figure 4.**
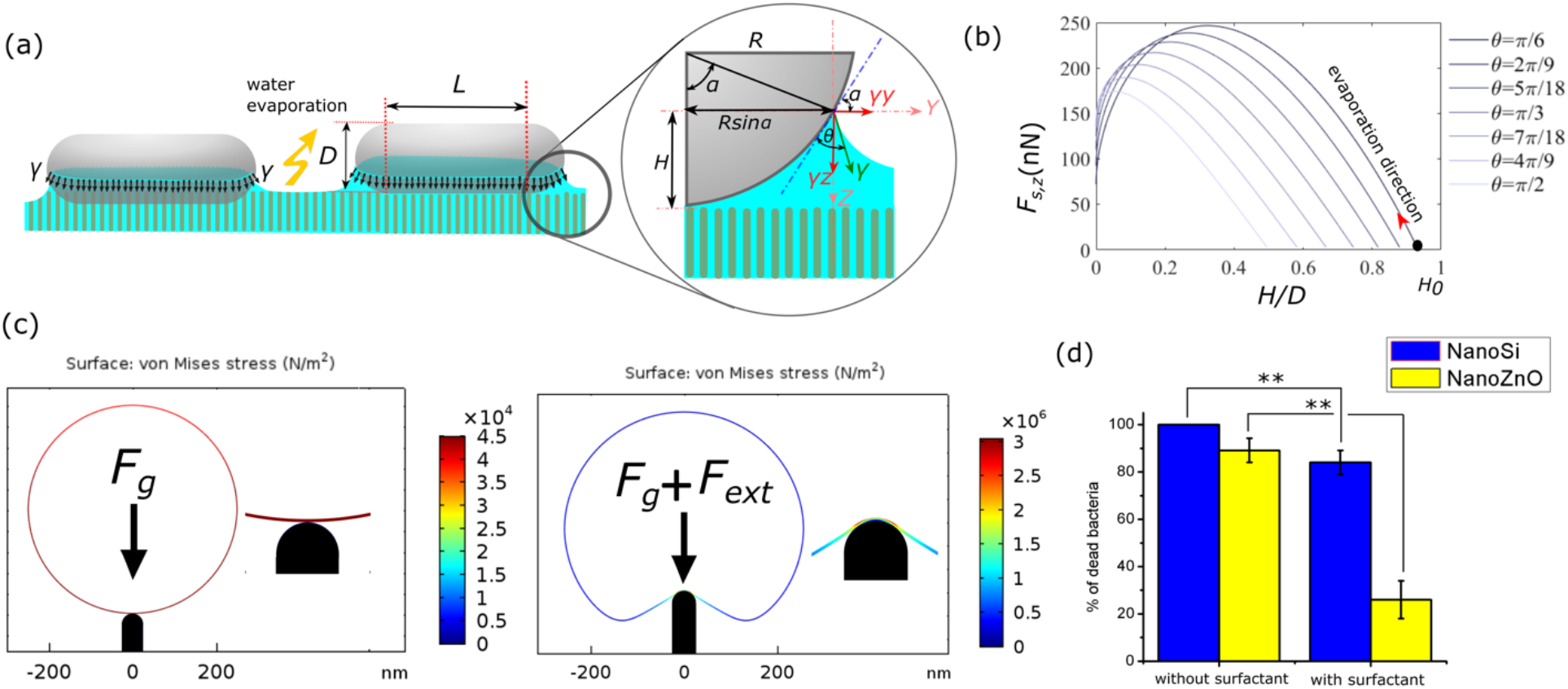
(a) Sketch of the capillary meniscus on bacterium during water evaporation and analysis of capillary forces; (b) the magnitude of the surface tension force as a function of bacterial contact angle (θ) and the liquid level (H) during water evaporation (H0 denotes the liquid level corresponding to initial equilibrium position of interface); (c) Numerical simulation of bacterial deformation under gravity (Fg) in the absence and presence of an external force (Fext) on NanoSi (refer to Figure S4 for NanoZnO). The plots show the von-Mises stress profile on bacterial cell envelope. The external force in the right graph was adjusted to give a maximum von-Mises stress of equal value to the cell wall ultimate tensile strength (for information about the simulation assumptions and parameters refer to SI); (d) Comparison of bacterial killing on NanoSi and NanoZnO in the absence and presence of Tween80. The viability of bacteria without surfactant is repeated from Figure 1b. Error bars denote the sample standard deviation and the experiment for each treatment was repeated three times. *Statistically significant difference (*p* < 0.05, n = 3, Student’s t-test).

Where *γ* is the surface tension and *C* is the perimeter of the contact line. Assuming a constant contact angle between the thin liquid film and bacterium during evaporation^35^, the integral of vertical component of the surface tension around the contact line yields the vertical force along the *z*-axis (*F_s_,_z_*). Assuming an idealized cylindrical bacterium geometry with two hemispherical caps, the magnitude of this force is:

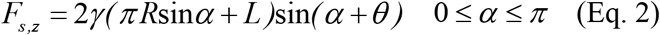

Where *L* is the length of the cylindrical part of the bacterium, *R* is the radius of the hemispherical edge of the bacterium, *θ* is the contact angle of the liquid with the bacterial surface, and *α* is the falling angle describing the position of three-phase contact line on the hemispherical end of a bacterium. The second component of normal capillary forces is caused by pressure differentials across the curved liquid/air interface. We experimentally determined that both surfaces were strongly hydrophilic, with water droplet contact angles of 5° and 20° respectively, for NanoSi and NanoZnO. Since these values are low, the droplet interface is nearly flat, and hence contributes only a small pressure differential across the interface. Wicking of liquid between nanopillars ensures that this condition remains valid during the evaporation timecourse^36^. As such, although capillary pressure tends to further increase the total downward capillary force, we assume that this effect is negligible^37^ under these operating conditions. Substituting *L*~1.0 μm, *R*~250 nm, *γ*=72 mN/m^38^ in Eq. 2, *F_s_,_z_* was calculated for *α_0_<α <*π where *α_0_* is the initial equilibrium position based on the flat interface assumption and π/6*<θ<*π/2 which covers the range of bacterial surface contact angles reported for *P. aeruginosa*^39–41^. For each value of *θ*, *F_s_,_z_* increases as water evaporates (*H*, liquid level, decreases), reaches a maximum value, and then decreases (Figure 4b). The maximum value of *F_s_,_z_* is between 150 nN and 250 nN for the above range of contact angles.

To determine whether these forces are sufficient to stress the bacterial cell wall to the point of rupture, we simulated the cell wall in contact with a nanopillar under gravity and an additional “external” load to capture surface tension forces in a lumped parameter model (Figure 4c). Under gravity alone, the von-Mises stress is significantly smaller than the cell wall tensile strength, and additional normal forces of ~0.1 nN for NanoSi and ~ 1 nN for NanoZnO are required to exceed the cell envelope ultimate tensile strength (Figure S7). The normal capillary forces at the air-liquid interface approach ~35× this amount for NanoSi and ~15× for NanoZnO (Figure S8; assuming that ~20 NanoSi pillars and ~6 NanoZnO pillars are in contact with each cell, with equal reaction forces from each of the pillars). Hence, the external forces generated by the air-liquid interface are theoretically able to rupture bacterial cells on nanopillars.

This computational model has intrinsic limitations, including an inability to capture nanopillar or cell wall heterogeneity, or complexity of the cell-surface interaction. Surface tension is a critical component of the model as the capillary force is linearly dependent on the magnitude of surface tension. Therefore, to experimentally validate its importance, we manipulated surface tension properties by adding a surfactant to the test solution. Tween80 surfactant at 0.5 wt% concentration reduces the water surface tension by approximately 65%, and is nontoxic to *P. aeruginosa* at this concentration in submerged conditions (data not shown). Adding surfactant significantly reduced the antibacterial efficacy of the nanopillars upon drying by ~74% on NanoZnO and ~20% on NanoSi (Figure 4d). Hence, reducing the surface tension and consequently the applied normal forces during air-liquid interface movement rescues bacterial viability, demonstrating that a critical external load is required for mechano-bactericidal activity.

To confirm that normal forces specifically are responsible for the bactericidal phenomenon, we next applied external compressive loads to directly deliver normal forces to bacterial cells on the nanopillars. A weighted 3D-printed stamp featuring an array of circular posts was pressed against NanoSi or NanoZnO surfaces while immersed in the bacterial suspension (Figure 5a). The apparent stress (σ) delivered to each surface was calculated based on the weight used to load the stamps against the surface. On flat control samples, all cells remained viable at the highest applied stress in this experiment. For areas of the NanoSi or NanoZnO surfaces that made contact with the stamp’s posts, circular regions of dead bacteria were produced for sufficiently high applied stresses while no cell death was observed in the regions between the compressive pillars (Figure 5b,c). Stresses as low as 0.17 MPa can produce a kill region on NanoSi but not on NanoZnO, which required a larger stress (0.4 MPa and above) to achieve the same antibacterial effect (Figure 5d). This correlates well with our earlier observations demonstrating that topography alone cannot drive bacterial death, but that a coupled external force with the nanopillar surface is necessary to deliver critical cell damage.

**Figure 5.**
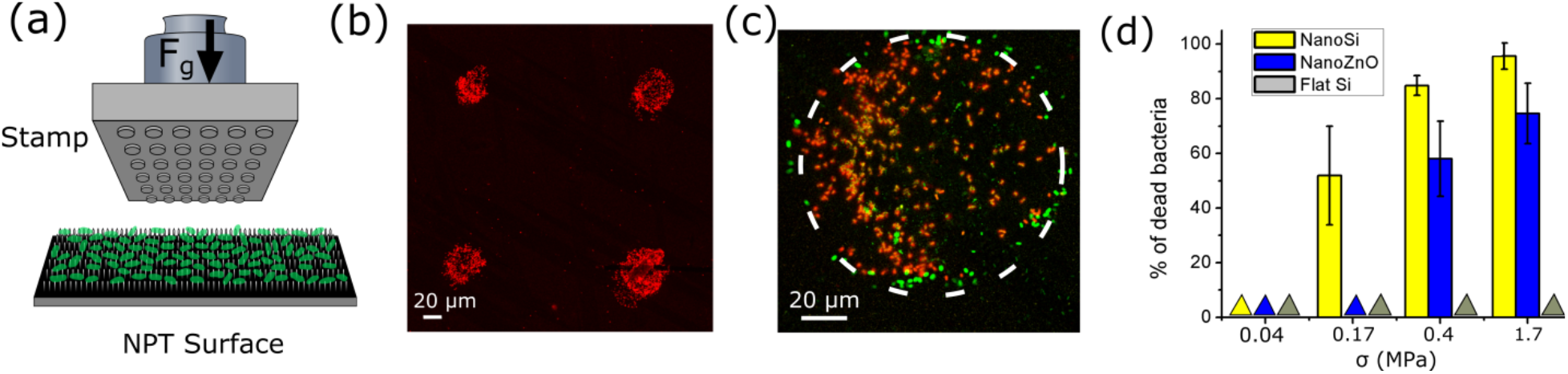
Mechanical compression test: (a) sketch of an external gravitational force (F_g_) applied by a 3D-printed stamp on bacteria attached to nanopillars; (b) kill-zone layout on NanoSi after the stamp contact at σ = 0.4 MPa; (c) dead and live bacteria in the kill-zone on NanoSi at σ = 0.4 MPa. The kill zone resembles the circular areas of microposts on the stamp (dashed line) (d) percentage of killed cells as a function of σ. Triangles represent the detection of no kill zone (i.e., all bacteria were viable). Error bars denote the standard deviations and experiments were repeated three times.

Previous researchers have suggested several different forces potentially responsible for damaging and rupturing bacterial cells on nanopillars. The most highly cited explanation of the mechano-bactericidal mechanism, by Pogodin *et al*.^8^, attributes the bactericidal activity to adhesion forces passively generated by the attachment of cells on nanopillars, which, in turn, stretch and rupture the cell membrane. This has been challenged by others who argued that in many cases, adhesion forces are not of sufficient magnitude to kill bacteria; thereby proposing additional contributions of other external forces^23, 42–43^. Other forces suggested to date include gravity^42^, motility forces^23^, and released bending energy from flexible nanopillars^25^. In our study, we encountered an experimental condition where none of the above forces seems to be dominant – as evidenced by the absence of bactericidal effect in fully immersed conditions. We also showed that alternative sources of normal external forces such as passage of an air-liquid interface or direct application of force can be employed to induce bactericidal responses in submerged conditions.

The dominant effect of capillary and mechanical forces on the bactericidal properties of nanopillars observed in this study has important implications in interpreting the results from other mechano-bactericidal experiments. For instance, considering the fast rate of evaporation of water on nanopillar surfaces (Figure S9), capillary forces can quickly arise in the time between sample preparation and fluorescence microscopy imaging. This experimental artefact can also occur by omitting CPD before SEM. Given the importance of these forces, we suspect that lack of attention to these forces might have partly contributed to the conflicting data on mechano-bactericidal mechanisms.

The significant role of external forces also reveals new insights into the practical utility of bactericidal nanopillars. Natural capillary forces that occur upon water evaporation may hence be desirable, and nanopillar surfaces can be designed for use in situations involving intermittent exposure to bacterial contamination through deposited droplets. surfaces in healthcare environments that contribute significantly to the spread of hospital-acquired infections may be designed with this use profile in mind. Alternatively, for applications requiring antibacterial properties in submerged conditions, such as anti-biofouling materials in industry or medical implants, we suggest that a controllable external force may be required to initiate/enhance mechano-bactericidal action. Designing systems to create suitable external forces in conjunction with nanopillar structures should hence be a fruitful and important research topic in designing topographical materials for practical antibacterial activity.

In summary, we fabricated two nanopillar surfaces, NanoSi and NanoZnO, and exposed them to the bacterium *P. aeruginosa*. Both surfaces showed no antibacterial activity when submerged in bacterial suspension for one hour. However, small droplets of bacterial suspension dispensed onto the surfaces quickly experienced cell death upon water evaporation. Real-time fluorescent microscopy, CPD-SEM, and computational simulations reveal that capillary forces that arise during droplet evaporation are responsible for rapidly killing *99.9%* of bacteria on NanoSi and 90% of bacteria on NanoZnO. We further demonstrated that a controlled mechanical force can efficiently kill bacteria on nanopillars under fully immersed conditions. These findings establish that an external force is required to effectively achieve bactericidal action on nanopillars.

## Supporting information

Supplementary figures and experimental details

## Acknowledgements

This research was supported, in part, by the Natural Sciences and Engineering Research Council of Canada (NSERC) and the McGill Interdisciplinary Initiative in Infection and Immunity (MI4) funding (NT, DN). We acknowledge the support of the Canada Foundation for Innovation, the Canada Research Chairs program (NT, CM), and the McGill Engineering Doctoral Award (AV and NL). We thank Reghan J. Hill for useful discussions. We are also grateful to the Advanced Bioimaging Facility and the Facility for Electron Microscopy Research at McGill University and CMC Microsystems for access to simulation software and to the Montreal Insectarium for providing cicada samples.

