## Supplementary figures and experimental details for "Mechano-bactericidal nanopillars require external forces to effectively kill bacteria"

### Supplementary Information (SI)

#### Materials and Methods

##### *NanoSi fabrication*

NanoSi substrates were fabricated based on methods outlined by *Peng et al.* [1]. Briefly, silicon wafers were cleaned in acetone, then isopropanol, rinsed with de-ionized water three times then soaked in Nano-strip solution (VWR, USA) at 65 ºC for 20 min. They were then rinsed thoroughly with de-ionized water before being dipped in buffered oxide etchant (Sigma–Aldrich, USA) for 1 min. Silver nanoparticles were electroplated onto the cleaned silicon wafer by soaking the surface in a solution of $AgNO_3$/HF ($AgNO_3$: 0.01 M, HF: 4.6 M) for 1 min. After that, the substrates were soaked for etching in a solution of $FeNO_3$/HF ($FeNO_3$: 0.135 M, HF: 4.6 M) for 45 min at room temperature. The obtained samples were rinsed again with de-ionized water and dried at room temperature.

##### *NanoZnO fabrication*

Hydrothermal synthesis of NanoZnO is a two-step process. First, 1 mM of zinc acetate dihydrate dissolved in anhydrous ethanol was spin-coated (2000 rpm, 30 s) onto glass microscope slides which were pre-cleaned with isopropanol and acetone and dried with nitrogen flow. The glass slides were then heated on a hotplate at 300 °C for 10 min to anneal the zinc to form a seed layer. In the second step, the glass slides were immersed vertically in an aqueous growth solution containing equimolar (25 mM) zinc nitrate hexahydrate and hexamethylenetetramine and left in an oven at 70 °C. After 24 h of growth, slides were removed and rinsed three times with de-ionized water then dried at room temperature.

***Ordered nanopillar array fabrication***

Two highly ordered arrays of nanopillars were fabricated on a silicon wafer via e-beam lithography. In one array, the pillar-to-pillar spacing between adjacent nanopillars was 200 nm whereas the spacing in the other array was 400 nm. The height and diameter of both arrays were 300 nm and 80 nm, respectively. First, silicon wafers were cleaned as above and two resists were spin-coated on the sample, the primer hexamethyldisilazane followed by the negative e-beam resist Ma-N 2401. The average thickness was about 500 nm. Then, e-beam lithography was performed with an adaptive dose depending on diameter, density and periodicity of the final design with a modified Zeiss microscope (Leo 1540 XB). Following the exposure, the resist was developed by immersion in a solution of Ma-D 525. Next, the silicon surfaces were etched with an inductively coupled plasma (ICP) etching system at 20 °C in a mixture of sulfur hexafluoride and octafluorocyclobutane. The etching time was adjusted as a function of the nanopillar height (~100 nm/min). Finally, samples were cleaned with a solution of remover 1165 (Dow Chemical Company, USA) to remove the resists.

***Fluorescence assay using GFP-tagged bacteria***

To evaluate the activity of cells during water evaporation on nanopillars and flat surfaces, GFP-labeled *P. aeruginosa* was used. It has already been demonstrated that the decay in the expression of GFP correlates well with the loss in membrane integrity and cell death[2]. A particular feature of this technique is that, contrary to live/dead assay, it does not depend on the penetration of live/dead stains to indicate the loss in viability and as such is potemtially useful in studying bacterial viability upon drying. The first step in this test is to prepare a bacterial suspension as above. A droplet of the suspension was then placed on the nanopillar-textured or control surface, and the interaction of bacteria with the surface was monitored using a fluorescence microscope (Olympus IX71, Japan). The experiments were repeated three times for each sample. For better visibility of cells, images were converted to greyscale in ImageJ (National Institute of Health, USA) and contrast and brightness were adjusted in the images all together.

***Computational simulations***

Numerical simulations of a bacterium deformation on a nanopillar were performed using structural mechanics module of COMSOL Multiphysics v.5.3 (Comsol Inc., USA) in 2D-axisymmetric geometry. The bacterium was modeled as a thin elastic shell representing the cell envelope (thickness=2.5 nm[3]) with a constant internal volume to account for the incompressibility of the internal fluid. A contact boundary condition between the lower surface of the shell and the nanopillar was specified and a prescribed vertical displacement towards the nanopillar was defined for the rest of the shell external surfacs. Also a constant pressure boundary condition was assigned to the internal surfaces of bacterial shell. In preparing the model, for simplicity, we ignored the effect of friction and assumed a linear elastic material for the bacterial cell envelope. Also the weight of the bacteria was specifed as a constant body force acting on the entire bacterium. Stress profiles and pillar reaction forces were plotted for bacterium displacement increments. In preparing the model, we assumed a Young's modulus of 10 MPa and ultimate tensile strength of 3 MPa for the cell wall, based on published estimates[4], and an internal hydrostatic pressure up to $10^5$ Pa, a typical value for bacterial turgor pressure[5].

***Transmission electron microscopy (TEM)***

The nanopillars of as-synthesized NanoSi and NanoZnO surfaces were individually observed by TEM (FEI Tecnai 12 BioTwin 120 kV TEM, FEI Company, USA) after scratching the nanopillar surfaces with a razor blade.

***Contact angle measurements***

Static contact angles of NanoSi and NanoZnO were measured on an OCA 20 contact angle analyzer (DataPhysics Instruments, Germany) equipped with automated drop delivery and digital camera. The measurements were performed in ambient temperature using the sessile drop method. A 3 μL water droplet was dispensed on the surface of each sample and the contact angle was measured after 3 s. All experiments were repeated three times and the average value of contatct angle was reported.

| | Diameter (nm) | Height (nm) | Spacing (nm) |
|---|---|---|---|
| NanoSi | 50±10 | ~900 | 447±395 |
| NanoZnO | 230±65 | ~1200 | 408±392 |

**Table S1**. NanoSi and NanoZnO nanopillar dimesions

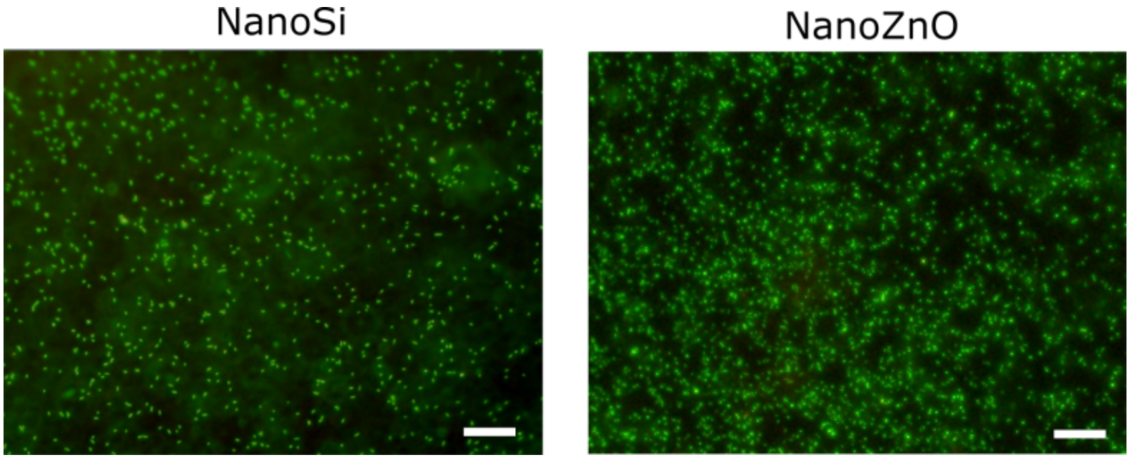


**Figure S1.** Viability of bacteria on NanoSi and NanoZnO after 1 h in the fully wet condition. The majority of cells retained green fluorescence and did not uptake red fluorescence, indicating they remain alive (scale bar = 20 μm).

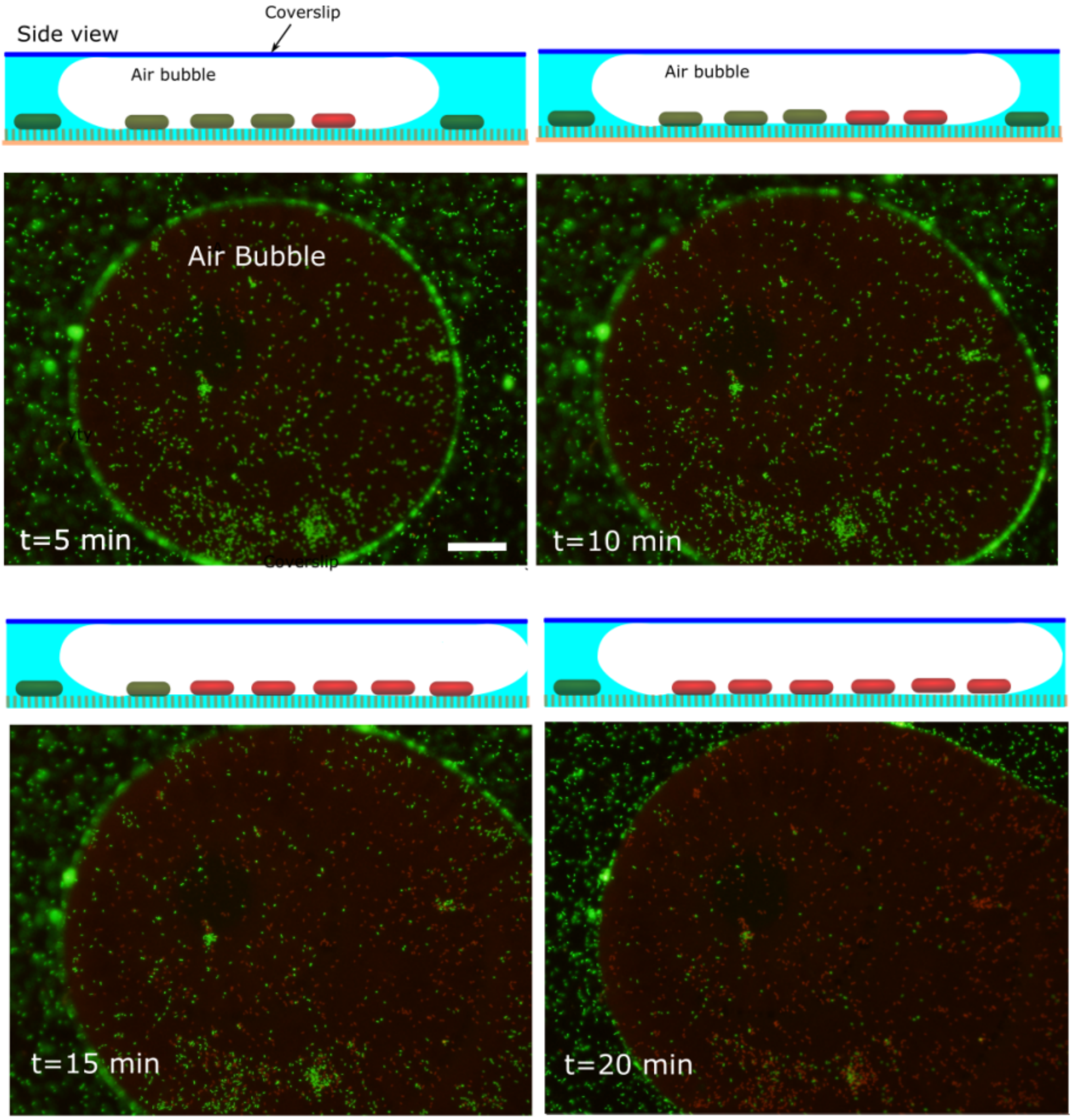


**Figure S2.** Viability of bacteria after the formation of an air bubble inside the confined space between NanoSi and a coverslip (scale bar = 50 µm). Top schematics above each image shows a side view of the possible bacterial killing process in a microbubble in the confined space. More than 50% of the population loses its viability in 5 min and more than 99% within 20 min of microbubble formation. The bacterial killing inside the microbubble is caused by capillary force due to the formation of air/liquid interface and subsequent liquid evaporation. It must be noted that the rate of water evaporation in this case is substantially smaller than a surface freely exposed to ambient due to the relatively small volume of air in the microbubble.

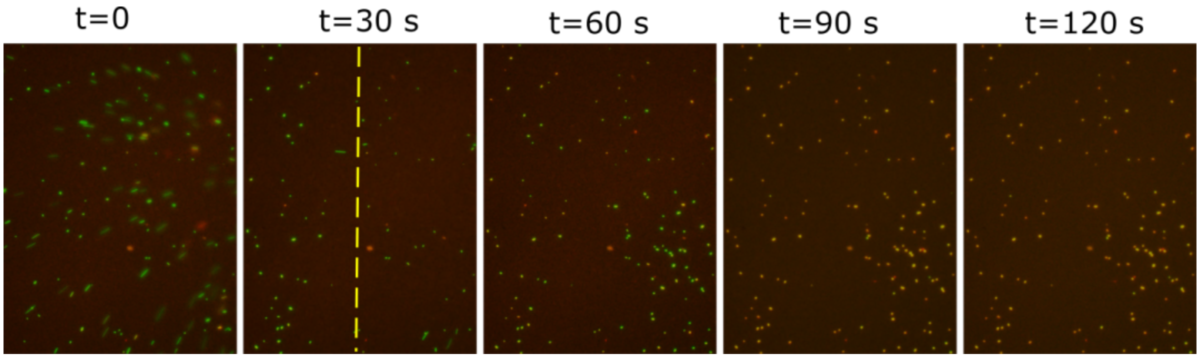


**Figure S3.** Viability of bacteria with time on NanoZnO upon passage of air/liquid interface (yellow line) during water evaporation (scale bar = 20 μm).

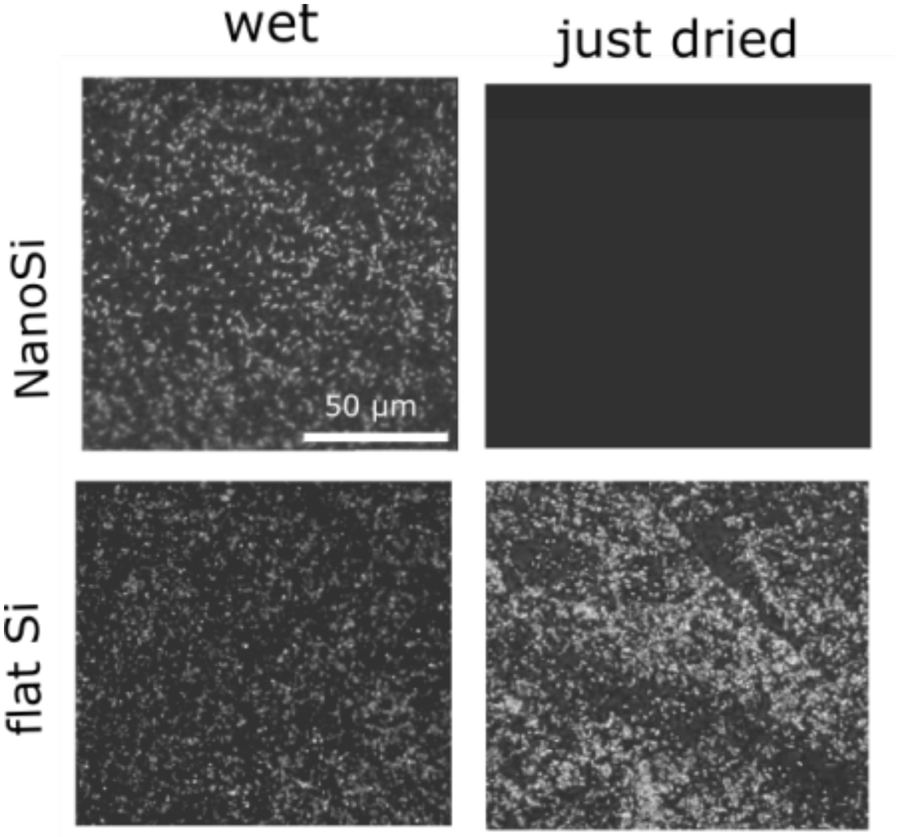


**Figure S4.** Fluorescent images of GFP-tagged *P. aeruginosa* on NanoSi and control (flat) Si in the wet condition and just after water evaporation. On wet NanoSi and NanoZnO surfaces, bacteria fluoresce green however, the bacterial fluorescence almost entirely vanishes after evaporation of liquid on the surface. On the control surface, attached and suspended bacteria fluoresced green in wet condition and remained fluorescent after liquid evaporation. According to Lowder *et al.*[2], the loss of bacterial GFP signal is indicative of loss in viability and could signify severe damage to cell membrane. The disappearance of GFP-signal, which is integral to cells, confirms that cell death observed in the live/dead assay is not caused by any artifacts relating to stains. In combination, both the live/dead and GFP assays demonstrate a rapid bactericidal mechanism on nanopillar surfaces upon water evaporation.

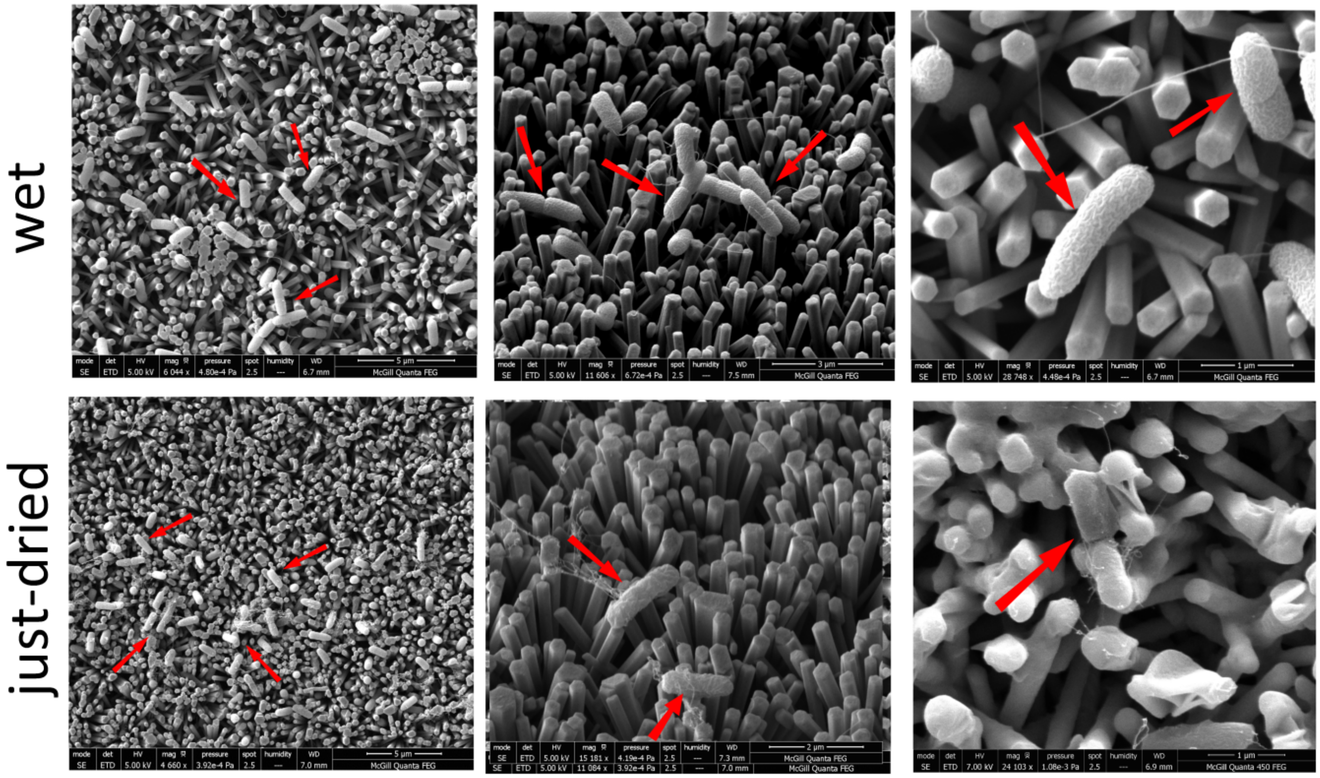


**Figure S5.** SEM of bacteria on NanoZnO in the wet condition and immediately after the liquid evaporation. Red arrows show the location of bacteria.

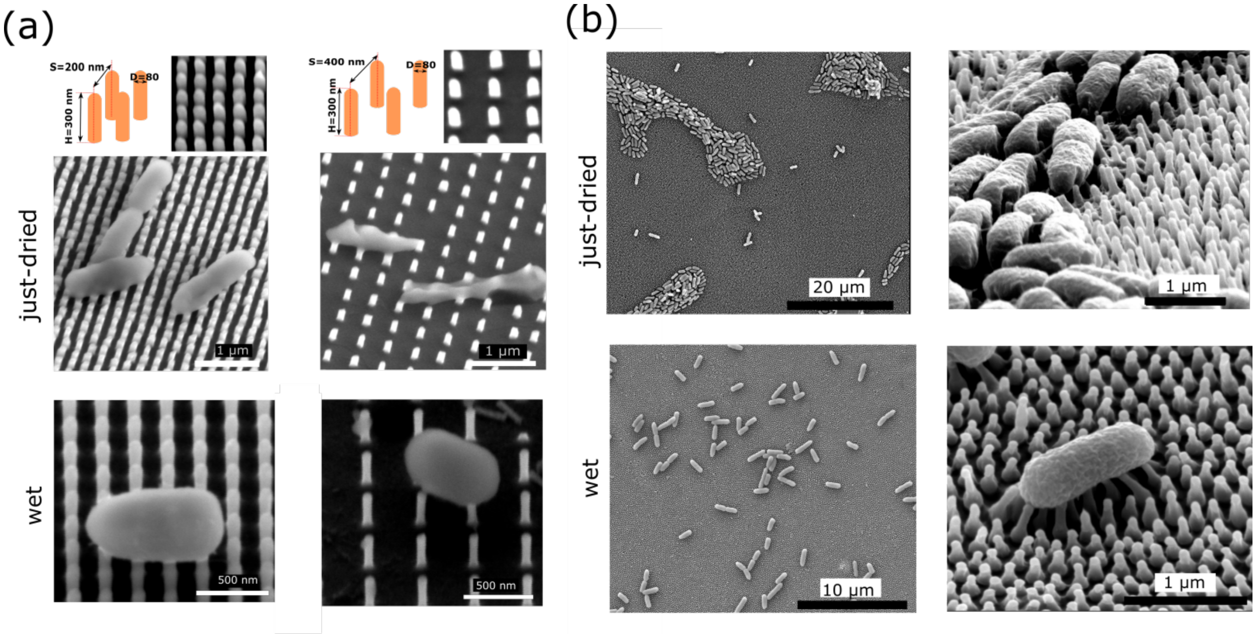


**Figure S6.** SEM images of bacterial morphology in the immersed wet state, where surfaces are not exposed to air during sample preparation, and the just-dried state on (a) two different ordered nanopillar arrays **(**dimension of each shown above the images). The extent of damage is greater on the pattern with larger spacing between the pillars. This is consistent with the theory of external forces. As the pillar array becomes denser the reaction to the external capillary force get smaller leading to less severe damage; (b) cicada *Salvazana mirabili*s (diameter = 90 nm, height = 300 nm, center to center spacing = 270 nm). We observe that bacterial structures are intact in the wet state for all the surfaces but damaged immediately after water evaporation. On cicada wings, we observe a difference in the distribution of bacteria on the wing surface between a wet and a dried state. Given that the wing surface is hydrophobic, bacteria tend to form agglomerates as a result of surface tension pulling the bacteria together in evaporating droplets. We did not pursue a comprehensive live/dead analysis for these surfaces due to limitation in sample supply for cicada wings and small pattern area (200 µm × 200 µm) for ordered nanopillar arrays.

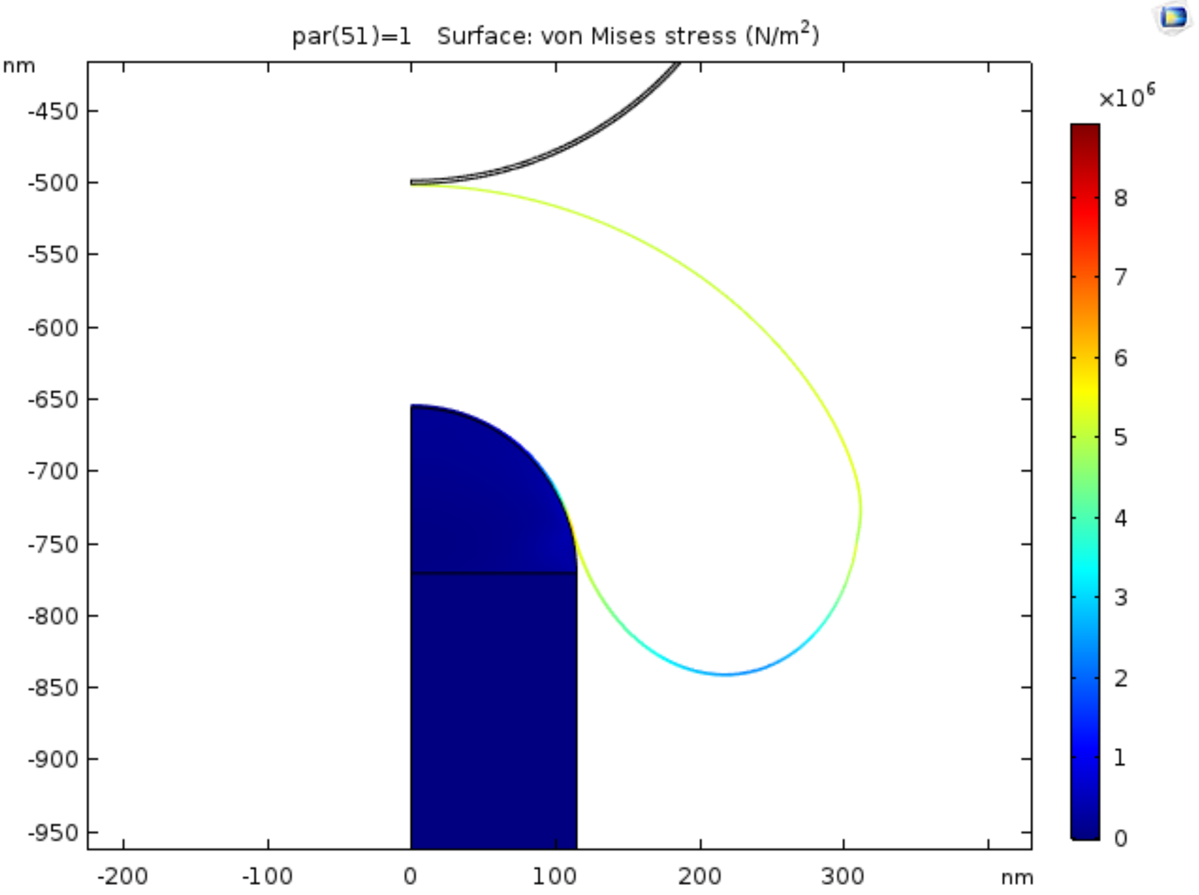


**Figure S7.** Stress profile of a bacterial envelope on a NanoZnO pillar. The force required to achieve the maximum von-Mises stress of 3 MPa (equal to the cell envelope ultimate tensile strength) is $1.7\times10^{-9}$ N, higher than the calculated value for NanoSi in Figure 4. Following parameters were used in the simulation: cell wall thickness = 2.5 nm[3], Young's modulus = 10 MPa[4], ultimate tensile stress 3 MPa[4], bacterial turgor pressure $10^5$ Pa[5] (for more information on the simulation refer to SI methods) .

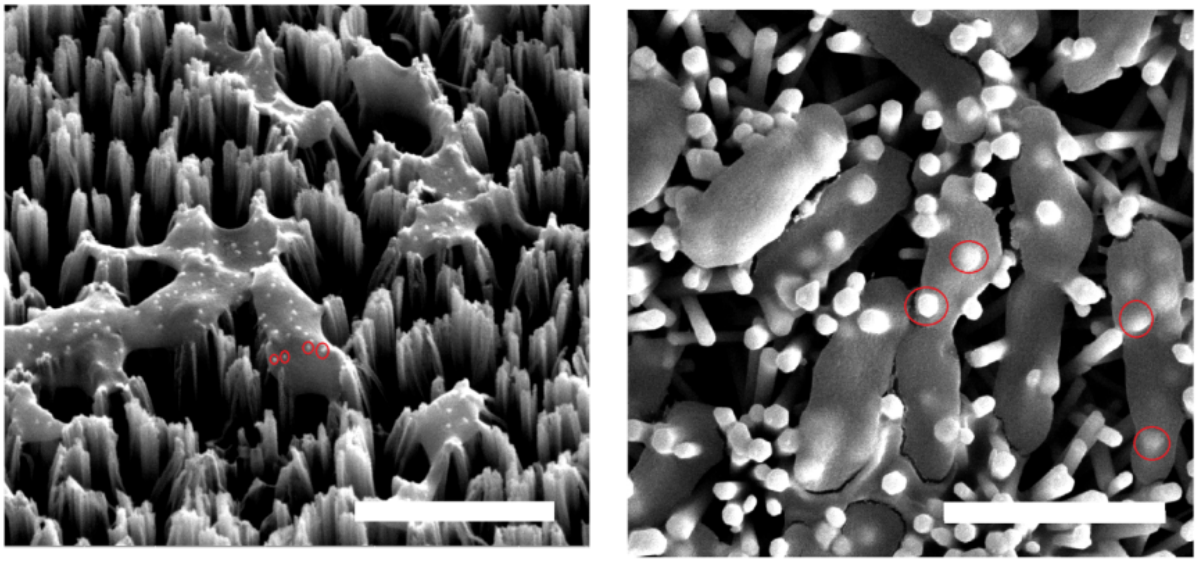

**Figure S8.** SEM images for estimating the number of nanopillars in contact with each bacteria. Bacteria are uncoated to make it possible to see the nanopillars (marked with red circles) penetrating through cells (scale bar = 2µm).

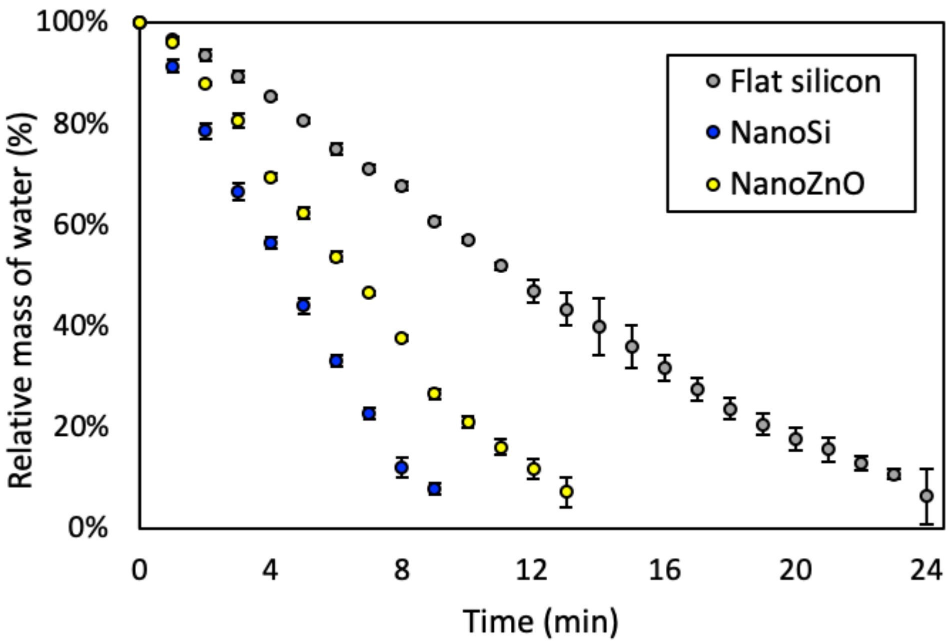


**Figure S9.** Droplets of water dispensed on NanoSi and NanoZnO evaporate 3× and 2× faster respectively compared to droplets dispensed on flat silicon since the hydrophilicity of the nanotopographies greatly increase the droplet spread area.
